# ZIKA - How Fast Does This Virus Mutate?

**DOI:** 10.1101/040303

**Authors:** Ian Logan

## Abstract

The World Health Organisation has declared the present epidemic of infection with the Zika virus to be a ‘Public Health Emergency of International Concern’. The virus appears to have spread from Thailand to French Polynesia in 2013, and has since infected over a million people in the countries of South and Central America. In most cases the infection is mild and transient, but the virus does appear to be strongly neurotropic and the presumptive cause of both birth defects in foetuses and Guillain-Barré syndrome in some adults.

In this paper the techniques and utilities developed in the study of mitochondrial DNA are applied to the Zika virus. As a result it is possible to show in a simple manner how a phylogenetic tree may be constructed and how the mutation rate of the virus can be measured.

The study shows the mutation rate to vary between 12 and 25 bases a year, in a viral genome of 10,272 bases. This rapid mutation rate will enable the geographic spread of the epidemic to be monitored easily and may also prove useful in assisting the identification of preventative measures that are working, and those which are not.

## INTRODUCTION

On the 1^st^ February 2016 the World Health Organisation declared the emerging Zika epidemic to be a ‘Public Health Emergency of International Concern’ which shows how this epidemic is now considered to be a major threat to the whole world (WHO, 2016). Their statement of intent includes the lines:

*Appropriate research and development efforts should be intensified for Zika virus vaccines, therapeutics and diagnostics*.

*National authorities should ensure the rapid and timely reporting and sharing of information of public health importance relevant to this PHEIC*.

In consequence it can be expected that many research institutions will increase their studies into Zika and related viruses and many new scientific papers will appear in the coming months. At the same time it is to be expected that many more RNA sequences of the virus will appear in the public domain.

Also, as a result of the rapidly increasing importance of the Zika virus, it is likely that scientists and physicians who normally would not consider studying the genetics of a virus might start looking at the newly available data.

The Zika virus is a *Flavivirus* carried by mosquitoes and was originally found in a Rhesus monkey identified in the Zika forest of Uganda in 1947, as described by Haddow et al. (1964). Over the next 60 years it was the cause of epidemics in several African countries. However, in about 2010 the virus spread to parts of Asia, in particular to Thailand (Fonseca et al., 2014; Haddow et al., 2012), and by 2013 had reached French Polynesia (Baronti et al., 2014). Since then there has been an explosive epidemic affecting the populations of many countries in both South and Central America. At the present time this epidemic shows no signs of abating.

In the many small epidemics there was no indication of the virus causing anything but mild and transient infections. But in the recent epidemic in Polynesia cases of central nervous system damage were observed and described as being a form of Guillain-Barré disease (Korff, 2013; Winer, 2014). However, in the current Brazilian epidemic the emphasis has been on the possibility of an association with birth defects, especially microcephaly resulting from maternal infection with Zika in the first and second trimester of pregnancy (Mlakar et al., 2016). The presumptive link between Zika infection and microcephaly is now looking more and more likely. Further cases of Guillain-Barré syndrome have also been seen.

The Zika virus is closely related to the viruses of Yellow Fever, Dengue and West Nile Fever, all of which do cause significant illness and mortality. However there are many other flaviviruses that are less well known and their hosts include horses, sheep, bats, birds and many other animals. A paper in 1998 listed over 70 different flaviviruses (Kuno et al., 1998) and new ones continue to be identified (Moureau et al., 2015).

All flaviviruses appear to have much the same structure as each other. The mature virus particles, virions, are about 50 nm in diameter and icosahedral in shape. Modern electron microscopy can now show the virion in considerable detail (Zhang et al., 2013; Zhou, 2014). The outer part is formed by an *envelope* overlying a phospholipid bi-layer *membrane* and the core contains a single stranded RNA molecule of about 10k bases.

In the mature Zika virion the RNA molecule, which encodes a *polyprotein* is described as having 10,272 bases, or 3,424 3-base codons for specific amino acids. The translation of bases to functional codons is not perfect, but for analysis purposes it has become accepted to describe the structure of the molecule in this manner: starting with MKN …. and ending with ….GVL (ie. The codons for: methionine, lysine, asparagine …. glycine, valine, leucine)

The *polyprotein* is a linear assembly of both *structural* and *non-structural* genes. The structural genes are for the *envelope, membrane and capsid*, and the non-structural genes are usually considered as being NS1, NS2A, NS2B, NS4A, NS4B, and NS5 (Bollati et al., 2010); and for the purposes of this paper this simple explanation will suffice.

The *envelope* and *membrane* genes define how the outer part of the virion is conformed. This outer part is important as it acts as an antigen for antigen-antibody reactions and also in the interaction between the virus and *entry receptors* as a virion attempts to enter a cell. In consequence, mutations affecting the construction of the *envelope* and *membrane* are probably more significant than mutations in other parts of the genome, and perhaps are of greater influence when it comes to possible changes in virulence.

It remains unclear as to which cells, if not all cells, in the human are susceptible to invasion with Zika. But the cells of the central nervous system do appear to be especially vulnerable (Mlakar et al., 2016) In all instances the process appears to be the same in that a virion attaches itself to the *entry receptors* on the outer surface of the cell and the virion enters the cell in a process described as endocytosis (Perera-Lecoin et al., 2014).

Once in a human cell the *envelope* and *membrane* separate from the core. The virus then hi-jacks the cellular apparatus for its own purposes. The *polyprotein* is copied and cleaved into its constituent parts and the daughter virions produced, each containing its own copy of the *polyprotein* (Clark & Harris, 2006).

At present there are few drugs that prevent replication of the Zika virus, and the older and well-established antiviral drugs, such as amantadine which are active against the influenza virus, are not helpful against the flaviviruses (Oxford et al., 1970). However a lot of work is being done to find new antiviral drugs (Sampath & Padmanabhan, 2009). It is interesting to note that a traditional Chinese remedy, Xiyangping, was used for its ant-viral properties in the treatment of a recent case of Zika (Deng et al., 2016).

In relation to the present epidemic the ability of the Zika virus to enter cells of the placenta and the central nervous system is particularly important. However for now it is unclear as to whether or not these invasions are more dependent on the strain of the virus, the genetic make-up of the host, or other factors. But once the virus has crossed the *placental barrier* to enter the foetus or the *blood-brain barrier* to enter the central nervous system it is likely that the usual antigen-antibody reactions are lessened and the virus is able to proliferate more easily. It is also unknown as to how long it might take for the virus to be cleared from the foetus or the central nervous system. Although it does seem likely that virus replication can continue in these areas for many months.

## METHODS

### The GenBank database

The RNA sequences for the Zika virus in the public domain can be found in the GenBank database of the National Institute of Health (Benson et al., 2013).

At present (March 2016) there are 16 complete sequences from virus collected in Africa and Asia before the start of the present epidemic and 17 sequences produced since.

Details of these sequences are given in Table 1.

**Table 1.** ZIKA RNA sequences on GenBank database (1 March 2016)

| Accession No. | Country of origin | Date of Collection |
| --- | --- | --- |
| African sequences |  |  |
| 1 LC002520 | Uganda | 1947 |
| 2 HQ234499 | Malaysia | 1966 |
| 3 HQ234500 | Nigeria | 1968 |
| 4 KF383116 | Senegal | 1968 |
| 5 KF383115 | CAR | 1968 |
| 6 HQ234501 | Senegal | 1984 |
| 7 KF268948 | CAR | 1976 |
| 8 KF268949 | CAR | 1980 |
| 9 KF268950 | CAR | 1980 |
| 10 KF383117 | Senegal | 1997 |
| 11 KF383118 | Senegal | 2001 |
| 12 KF383119 | Senegal | 2001 |
| Asian sequences |  |  |
| 13 EU545988 | Micronesia | 2007 |
| 14 JN860885 | Cambodia | 2010 |
| 15 KU681082 | Philippines | 2012 |
| 16 KU681081 | Thailand | 2014 |
| Current epidemic |  |  |
| Brazilian Reference Sequence |  |  |
| 17 KJ776791 | Polynesia | 2013 |
| 18 KU365779 | Brazil | 2015 |
| 19 KU365778 | Brazil | 2015 |
| 20 KU365777 | Brazil | 2015 |
| 21 KU365780 | Brazil | 2015 |
| 22 KU312312 | Suriname | 2015 |
| 23 KU501215 | Puerto Rico | 2015 |
| 24 KU509998 | Haiti | 2014 |
| 25 KU321639 | Brazil | 2015 |
| 26 KU527068 | Brazil | 2015 |
| 27 KU647676 | Martinique | 2015 |
| 28 KU501216 | Guatemala | 2015 |
| 29 KU501217 | Guatemala | 2015 |
| 30 KU707826 | Brazil | 2015 |
| 31 KU497555 | Brazil | 2015 |
| 32 KU740184 | China | 2016 |
| 33 KU744693 | Venezuela | 2016 |

The corresponding page on the GenBank database for a given sequence can be found by using a URL of the form:

http://www.ncbi.nlm.nih.gov/nuccore/KU744693

Each page give the amino acid list and the nucleotide base FASTA file for the RNA sequence. However, a GenBank page contains no real explanation as to what each list or file might mean and for this reason the author has developed a pair of *Zika virus utilities* that allows the user to compare one sequence with another.

### The Zika virus utilities

In conjunction with this paper 2 simple utilities have been prepared and are to be found the author’s website. These utilities are in the form of 2 webpages.

The URL for the pages is: www.ianlogan.co.uk/zikapages/zika.htm From where the user can choose to use: either the **Amino Acid Analyser** or the **Nucleotide Base Analyser**

The **Amino Acid Analyser** has in its source file copies of the amino acid lists for all the complete RNA sequences to be found in the GenBank database and a small Javascript program that allows the user to compare any sequence against any other. The results are displayed as a list of amino acid changes.

The **Nucleotide Base Analyser** has in its source file copies of the FASTA files for the complete RNA sequences of all the sequences for the current epidemic. Again, a small Javascript program that enables the user to compare two sequences and find the mutational differences between them.

Although the webpages can be viewed with any of the commonly used web browsers, the author recommends MOZILLA FIREFOX as this browser allows the user to alter the size of the text area, if needed.

It is the author’s intention to keep these webpages up-to-date as new Zika RNA sequences appear on the GenBank database.

## RESULTS

The results from this study can be considered under 3 headings:

### 1. Non-synonymous amino acid changes observed in the present epidemic

A mutation that causes a non-synonymous change of an amino acid is often considered to be significant. But if the change is between amino acids of similar size and of a similar polarity, there is probably no effective change in the functioning of the target protein.

The amino acid changes shown by the Zika RNA sequences in the present epidemic are listed in Table 2. The table demonstrates that by using this method the sequences can be split into 12 different strains with between 0 and 25 amino acid changes.

**Table 2.**
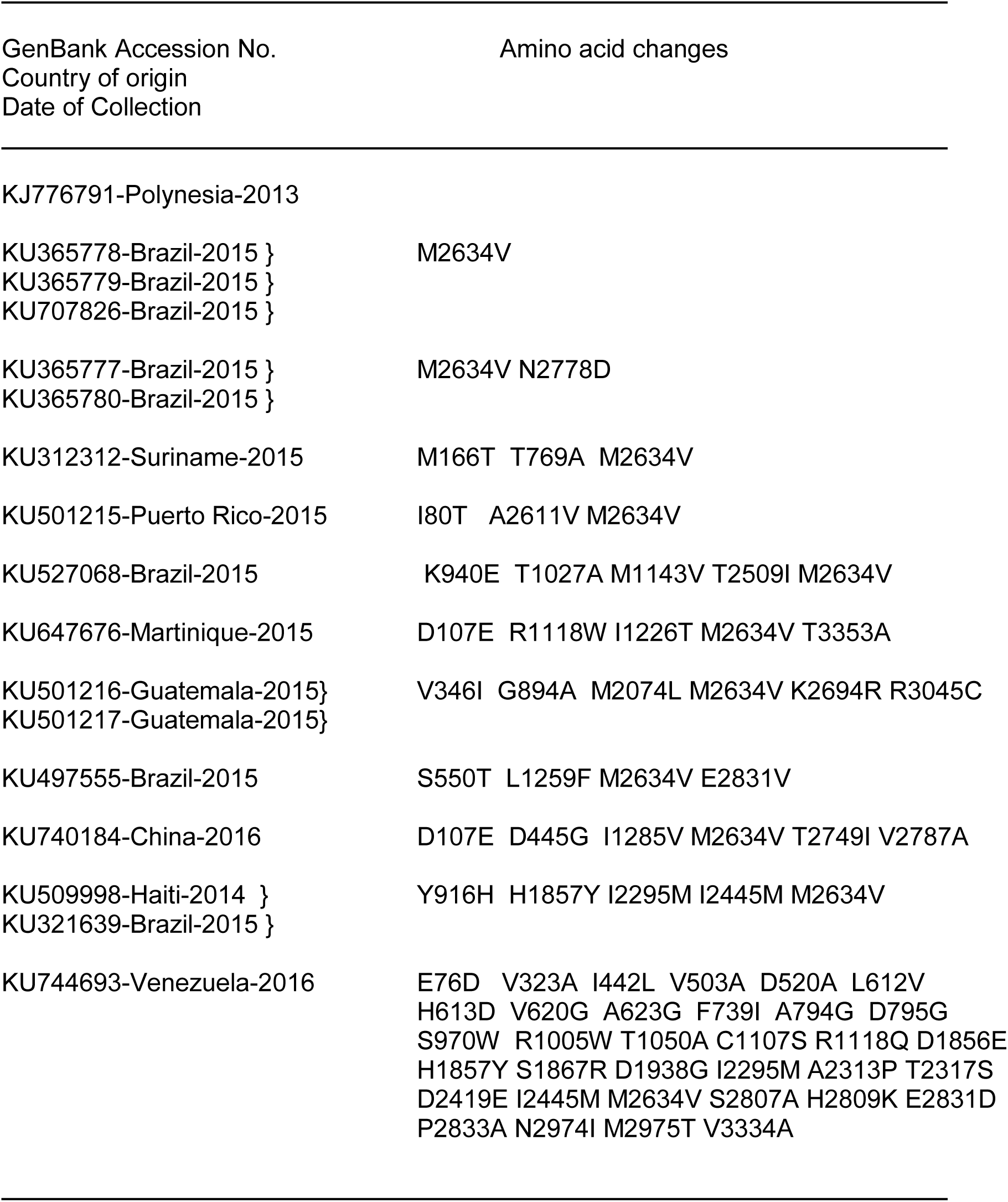
Non-synonymous amino acid changes found in sequences from the present epidemic. The change M2634V (Methionine to Valine), common to all the sequences occurs in the NS5 gene and is the result of the base mutation: A7900G which changes the codon from ‘ATG’ to ‘GTG’.

The mutation M2634V is common to all the virus strains that come from countries in South and Central America and is caused by the base mutation A7900G. However, as this mutation is found in the NS5 gene it is unlikely to be of significance as to the virulence or general behaviour of the Zika virus. The NS5 gene is involved in the replication of new virions and is not a structural gene (Zhao et al., 2015). It is perhaps too early to say that this mutation has absolutely no effect, but for the moment the M2634V mutation can be seen to be a useful marker to the present epidemic.

### 2. Base mutations in samples collected in the present epidemic

Whereas there are relatively few non-synonymous mutations in the virus strains collected in the present epidemic, there are many more synonymous mutations (i.e. mutations that do not produce a change of amino acid). This means that a meaningful phylogenetic tree can be constructed.

**Figure 1** shows the phylogenetic tree produced by using the mutations from the 17 complete Zika sequences currently to be found in the GenBank database. This figure shows the virus samples can now be separated into 15 different strains, considering the pairs of sequences KU365777/KU365780 and KU365799/KU707286 as being from 2 strains.

**Figure 1.**
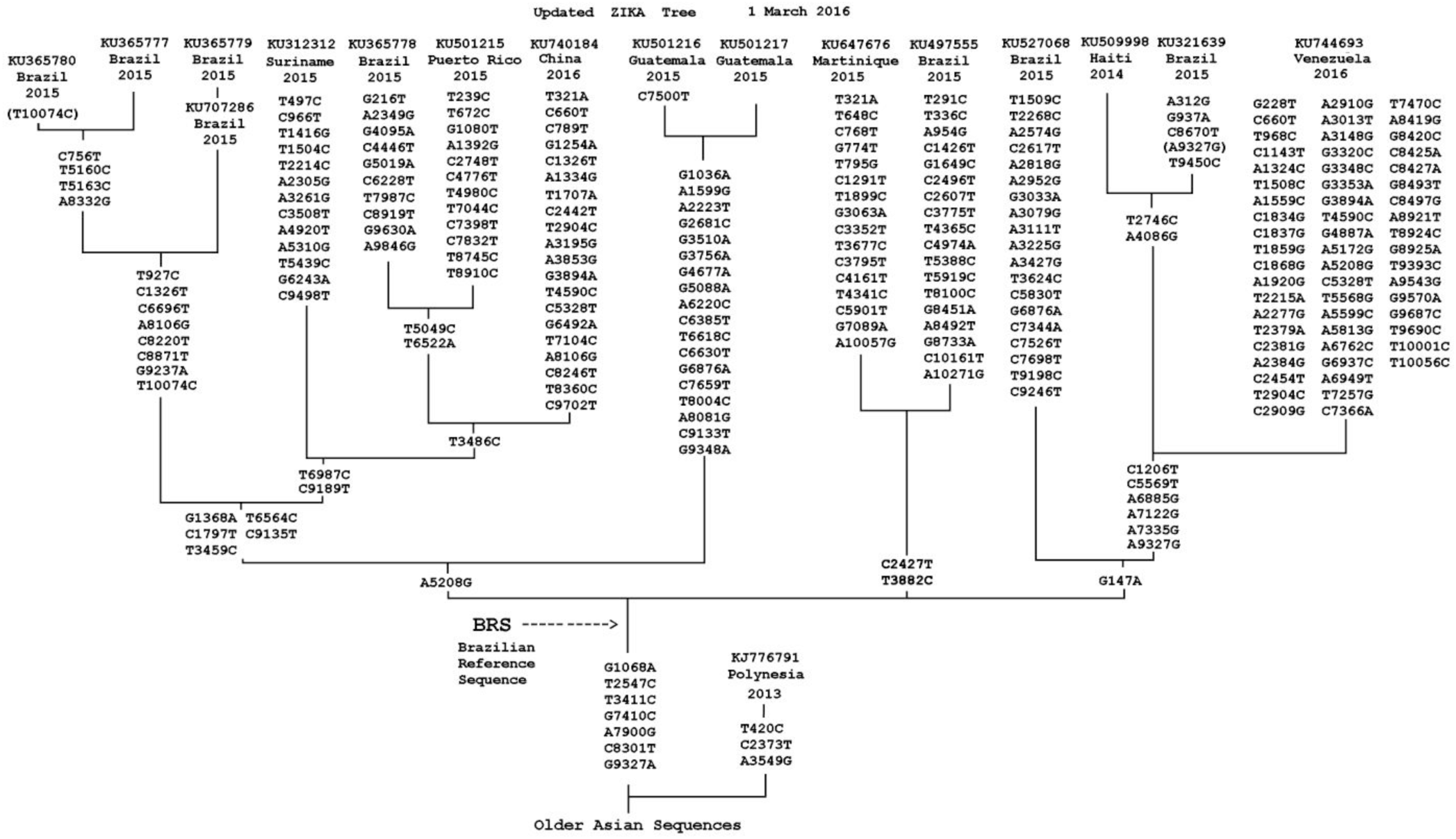
The phylogenetic tree of the 17 Zika RNA sequences from samples collected in the present epidemic. The 2 missing mutations - indicated by the brackets: C10074T in sequence KU365780-Brazil-2015, G9327A in sequence KU321639-Brazil-2015, are probably caused by technical errors. The position of a hypothetical *Brazilian Reference Sequence* (BRS) is marked on the tree. The BRS is used in the 2 utilities, the Amino Acid Analyser and the Nucleotide Base Analyser prepared for this paper and available at: www.ianlogan.co.uk/zikapages/zika.htm.

Note that in 2 of the samples there appear to be a *missing* mutation: G9327A in sequence KU321639-Brazil-2015 C10074T in sequence KU365780-Brazil-2015 and it is possible that these omissions result from technical errors.

### 3. Making an estimate of the mutation rate for the Zika virus

The data presented in **Figure 1** shows the actual mutations vary in number between 9 and 64 for the sequences from the strains of Zika virus collected in the course of the present epidemic. This number is calculated by considering the mutations that have occurred since the outbreak of the epidemic in Brazil and **Figure 1** suggests the use of a hypothetical *Brazilian Reference Sequence* (BRS) to describe a possible sequence for the original strain arriving in Brazil.

The number of mutations found in a sequence would appear to be proportional to the date on which the original sample was collected. The early sequences show between 9 and 30 mutations, whereas the two latest sequences, KU740184 and KU744693, show 30 and 64 mutations, respectively. The latter sequence is from a sample collected on the 6^th^ February 2016 and shows that the Zika virus continues to mutate at a rapid rate.

As the present epidemic can be considered to have started in Polynesia in 2013 (Baronti et al., 2014) and now to have lasted about 2.5 years (i.e. late 2013 to early 2016), the mutation rate appears to vary between about 12 to 25 mutations a year. The genome of the Zika virus is normally considered to be a *polyprotein* of 10,272 nucleotide bases, so the mutation rate can also be considered as changing between 0.12% and 0.25% of the RNA *polyprotein* each year.

It is not possible, using the data presently available, to give a more accurate value for the mutation rate. But the suggested rate of 12 to 25 mutations a year would appear to be a suitable starting point for further studies.

## DISCUSSION

### The Present Epidemic

The decision by the World Health Organisation to declare a *Public Health Emergency* in February 2016 because of the threat of a pandemic from Zika virus may be thought to have been a pessimistic move. However the evidence appears to indicate that the Zika virus is no longer restricted to localised habitats and largely dependent on the monkey as its host, but now covers a much larger area which includes many of the countries of South and Central America where it is wholly dependent on the human as its host. Unfortunately, there appears to be little if any herd immunity against the Zika virus in the populations of these countries, despite the closely related Dengue virus being very prevalent.

The sudden spread of Zika to South and Central America does not appear to have been due to any change in the mosquito vector or anything known to make the Zika virus itself more virulent. But it is rather from the fact that infected people are now able to fly rapidly from country to country, thereby spreading the disease extremely easily. This means there is little to stop the epidemic continuing to spread to further areas of population which also have a low level of herd immunity against the virus.

The absence of mosquitoes and the low incidence of person-to-person spread of the virus will probably mean the epidemic will not spread in the countries of the Southern and Northern latitudes. But from the evidence obtained so far it would appear likely the epidemic is only at its earliest stage and any suggestions as to what might happen remain speculative (Bewick et al., 2016).

### The Zika phylogentic tree

The 2 utilities prepared for this study show that many distinct strains of the Zika virus now exist, even though the present epidemic is less than 3 years old. When considering just the non-synonymous mutations in the RNA it is now possible to define 11 strains in the present epidemic. However, a more detailed examination looking at the actual mutations of the available sequences distinguishes 15 strains. As more data is made available it is to be expected the number of identifiable strains will increase. The phylogenetic tree as shown in **Figure 1** suggests that there is beginning to be a geographical spread of the associated virus strains, with distinct strains now coming from Martinique, Guatemala, Puerto Rico and Suriname, whilst Brazil continues to show a mixture of strains.

### An Estimate of the Zika Mutation Rate

The data used in building the phylogenetic tree can also be used to estimate the mutation rate of the Zika virus; and this would appear to vary between about 12 to 25 mutations per year, which is equivalent to about 0.12 - 0.25% of the RNA mutating each year. This rate is very high when compared to the Human DNA mutation rate, where a period of perhaps 250,000 years might be expected (Logan, 2015). But really it is not appropriate to compare RNA mutations against DNA chromosomal mutations as DNA replication is a self-correcting process, whereas RNA duplication is liable to many sorts of error.

However, the clear evidence of a high mutation rate in the Zika virus will allow for the present epidemic to be tracked in a fairly simple manner; and also it should be helpful in seeing where a local initiative of mosquito prevention is working and where it is not.

### The Complications of Zika infection

A particular feature of the present epidemic is the presumptive link to the high incidence of foetal abnormalities and cases of Guillain-Barré syndrome. These two complications appear to be caused in very different ways with the foetal abnormalities possibly being the result of direct infection of the foetus, whereas in cases of Guillain-Barré syndrome is possibly due to an exaggerated auto-immune response (Cao-Larmeau et al., 2016; Willison et al., 2016).

However, in the author’s opinion both of these conditions may result from the same underlying cause in which virus gets across the normally impenetrable *placental barrier* and the *blood-brain barrier*. How this happens is unclear, but, perhaps this is just a matter of a person getting a very high initial infection, possibly by having bites from physically large carrier mosquitoes, or maybe bites from several carrier mosquitoes in a very short period of time.

A study using the West Nile virus (Styer et al., 2007) showed that whilst most of the inoculum from a mosquito bite remains localised in the skin, there is always a significant initial viraemia. In this respect the recent report from Slovenia (Mlakar et al., 2016) shows the x-rays of an affected foetus having numerous calcifications in the placenta and brain. Whilst it is unproven it would seem possible that these lesions result from localised ‘viral plaque formation’ associated with an initial viraemic spread. A similar clinical picture is seen in tuberculosis. Although this disease is caused by a bacterium and not a virus, the resulting x-ray picture of localised calcifications is well recognised and is termed miliary tuberculosis (Khan et al., 2011; Yang et al., 2015).

It is also possible that the risk of developing complications of Zika may reflect the genetic differences between the sufferers and the general population. But at the present stage of our knowledge there is no indication of what particular differences might be important.

### Conclusion

This study shows in a simple way how sequencing data from samples of the Zika virus available in the public domain can be collected and analysed. Using this data it is possible to construct a phylogenetic tree and show that in the present epidemic there are already many identifiable strains of the virus. The data can also be used to show that the Zika virus has a high mutation rate.

This short paper raises as many questions as it tries to answer. The present epidemic is from the Zika virus, but Yellow fever cases are rising in Africa and Dengue affects millions of people each year. So further pandemics caused by flaviviruses, other than Zika, pose a continuing threat.

